# Return of an apex predator to a suburban preserve triggers a rapid trophic cascade

**DOI:** 10.1101/564294

**Authors:** Kevin Leempoel, Jordana Meyer, Trevor Hebert, Nicole Nova, Elizabeth A. Hadly

## Abstract

Absence of apex predators simplifies food chains, leading to trophic degradation of ecosystems and diminution of the services they provide^1^. However, most predators do not coexist well with humans, which has resulted in a decline of carnivores and functional ecosystems worldwide^2^. In some instances, cryptic carnivores manage to survive amidst human settlements, finding refuge in small biological islands surrounded by urban landscapes. In such a system, we used two non-invasive data collection methods (camera trapping and fecal sampling) to investigate the multiannual relationship between predators and prey, and between competitors, through analysis of: (1) relative abundance and detection probability of species over time, (2) causal interactions via empirical dynamic modeling, (3) diet, and (4) diel activity patterns. All approaches show concordance in the results: the natural return of an apex predator, the puma (*Puma concolor*), triggered a trophic cascade, affecting the abundance and behavior of its main prey, subordinate predators and other prey in the studied system. Our study demonstrates that trophic recovery can occur rapidly following the return of a top predator, even in small protected areas in increasingly urbanized landscapes.

## INTRODUCTION

Populations of apex predators are in global decline, resulting in a major trophic downgrading of functional ecosystems and ecosystem services^1^–^3^. Trophic cascades — the top-down effects predators have on food webs across multiple trophic levels — are especially relevant to management efforts to re-introduce predators and restore ecosystem function^4,5^. Beyond the common tri-trophic model (carnivore-herbivore-plants), apex predators also influence food webs through intermediary species (e.g., omnivores and mesopredators)^6^. Top predators control surges of mesopredator populations and thus decrease pressure on subordinate mesopredators and prey^7,8^, or they may directly predate or scare prey, changing their foraging behavior, location, and vigilance^4,9^. However, documenting a dynamic trophic cascade in real time is rare and most studies instead rely on short-term monitoring of indirect evidence, or comparisons of systems with or without an apex predator^4^.

In North America, trophic cascades caused by pumas have not attracted the same attention as those caused by wolves^10^, despite pumas being the most widely distributed carnivore in the western hemisphere^2^. To our knowledge, only three studies on puma-mediated trophic cascades have been published to date, all of which relate to their extirpation from studied ecosystems^11^–^13^. Pumas are known to be subordinate to grizzly bears (*Ursus arctos)*, wolves (*Canis lupus)* and jaguars (*Panthera onca)* where these rarer predators still occur^14^. Pumas have become the apex predator across the Americas, despite some regional extirpation and their fragmented distribution^15,16^. Moreover, pumas are affected by human activities and tend to avoid humans both in space (*e.g.*, a lower occupancy correlated with human density^14^) and time (*e.g.*, increased nocturnal activity in high-versus-low human densities^17^).

Here, we demonstrate that the natural increase in resident pumas in a small exurban preserve (≈ 5 km^2^) was responsible for a multi-tiered trophic cascade, affecting both the abundance and behavior of its main prey, the mule deer (*Odocoileus hemionus*), and its competitor, the coyote (*Canis latrans*), which in turn had downstream effects on subordinate predators and prey. We employed a suite of different approaches to reveal this finding and its underlying mechanisms: (1) relative abundance index (RAI) and detection probability inference from long-term camera-trapping efforts, (2) empirical dynamic modeling to infer causal interspecies relationships from RAI data, (3) diet analysis of predators from fecal samples, and (4) daily activity cycle analysis to study behavior.

## RESULTS

From 2010 to 2017, 176446 pictures were collected in Jasper Ridge Biological Preserve (JRBP; Stanford, CA) in a total of 39621 trap days, with 9 cameras starting in 2010 (7-year dataset) and 16 in 2012 (5-year dataset), the latter set covering the preserve more extensively. Wildlife was captured in 50% of the photos, 29% contained humans and 21% were blank. We extracted independent photographic events for 11 mid-large animal species, but chose to focus on 5 species hypothesized to be part of a food web: puma, mule deer, coyote, bobcat (*Lynx rufus*), and gray fox (*Urocyon cinereoargenteus*) (Table 1).

**Table 1.** Number of independent photographic events recorded per species in both 7-year and 5-year datasets. Species are ranked in decreasing order of number of events in the 5-year dataset. Last column indicates the number of scats sequenced per species.

|  | # of events<br>in 7-year dataset | # of events<br>in 5-year dataset | Scat samples<br>sequenced per species |
| --- | --- | --- | --- |
| **Mule deer** | 11595 | 11996 | NA |
| **Coyote** | 3190 | 2166 | 11 |
| **Bobcat** | 679 | 1140 | 29 |
| **Puma** | 436 | 1046 | 13 |
| **Gray fox** | 330 | 452 | 46 |
| **All 11 species** | 20928 | 25353 | 99 |

The RAI and detection probability time series showed a substantial increase in pumas within an 18-month interval, which then stabilized (time points T1-T2 in Figure 1, S1 and S2). During that period, the RAI of mule deer and coyote decreased, and both were relatively stable after three years (T3). During this shift, the RAI of gray foxes increased substantially. Bobcats on the other hand, kept a similar detection probability (Figure S2) and RAI in the 7-year dataset, but the latter differed substantially from the 5-year RAI curve. Such inconsistency can be explained by the biased spatial coverage of the 7-year dataset. We also looked at coyote group size over time and found that events involving more than one individual were frequent before 2012 and subsequently declined, such that almost all coyotes are now observed as individuals (Figure S4).

**Figure 1.**
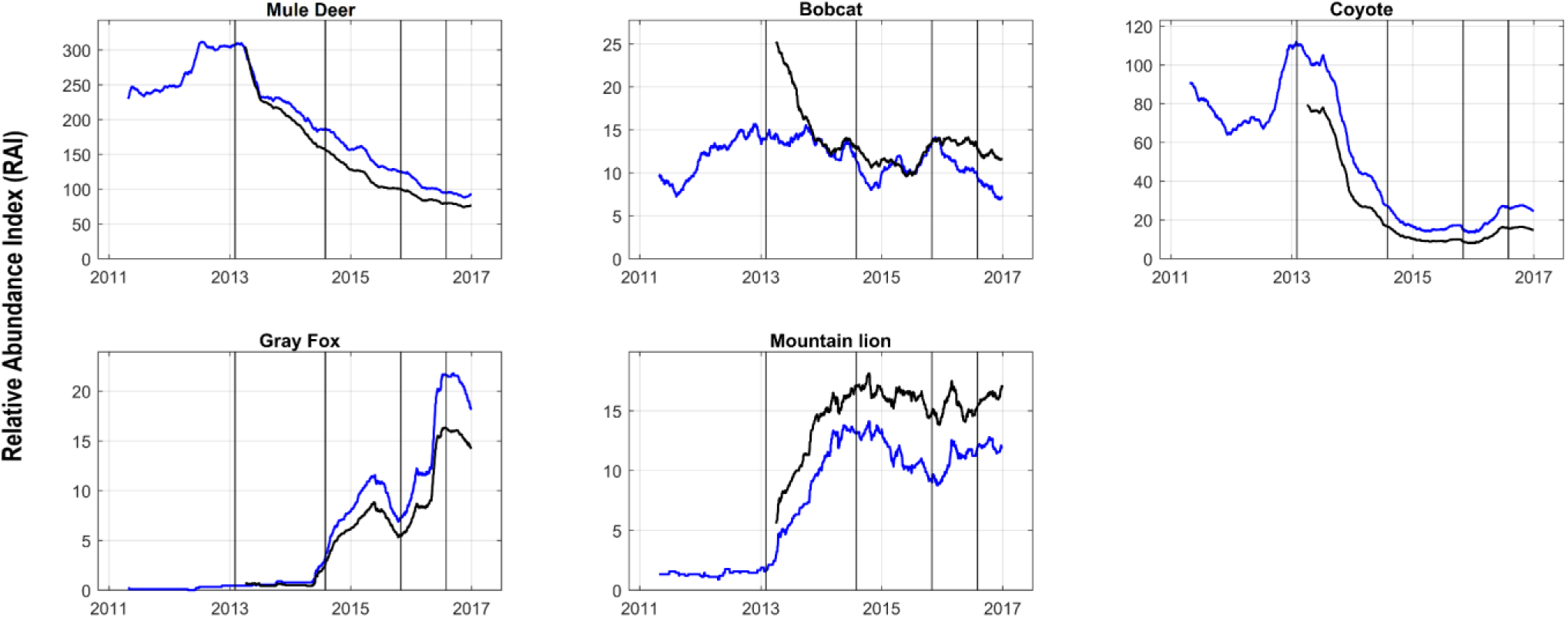
Relative Abundance Index (RAI) per year of the five species included in the 7-year (blue) and 5-year (black) datasets. Vertical lines correspond to selected time points T1 to T4: T1: 01-02-2013; T2: 01-08-2014; T3: 01-11-2015; T4: 01-08-2016.

In order to investigate causal relationships between the five species, we conducted an empirical dynamic modeling (EDM) approach called convergent cross-mapping (CCM)^18^ (Figure 2 and S3). This approach uses time series to reconstruct the dynamics of a system by constructing a state-space manifold using only the time series of the hypothesized response variable (*e.g*., deer RAI). This manifold is then used to infer the time series of the driver (*e.g*., puma RAI). If the inferred driver time series matches the observed driver time series (measured by cross-mapping skill and convergence), then CCM suggests that there is a causal relationship between the hypothesized driver and response variable (see Methods for details). The results primarily show that abundance of puma drives that of deer, coyote and gray fox (Figure 2a, 2b and 2d). There is some evidence of bottom-up feedback as well, although generally top-down regulation is stronger (higher cross-mapping skill). There seems to be a causal relationship between the canids as well (Figure 2e); however, the results from the 5-year dataset were not significant (Figure S3e). There is no significant relationship between bobcats and puma, and bobcats and coyotes (Figure 2c and 2f).

**Figure 2.**
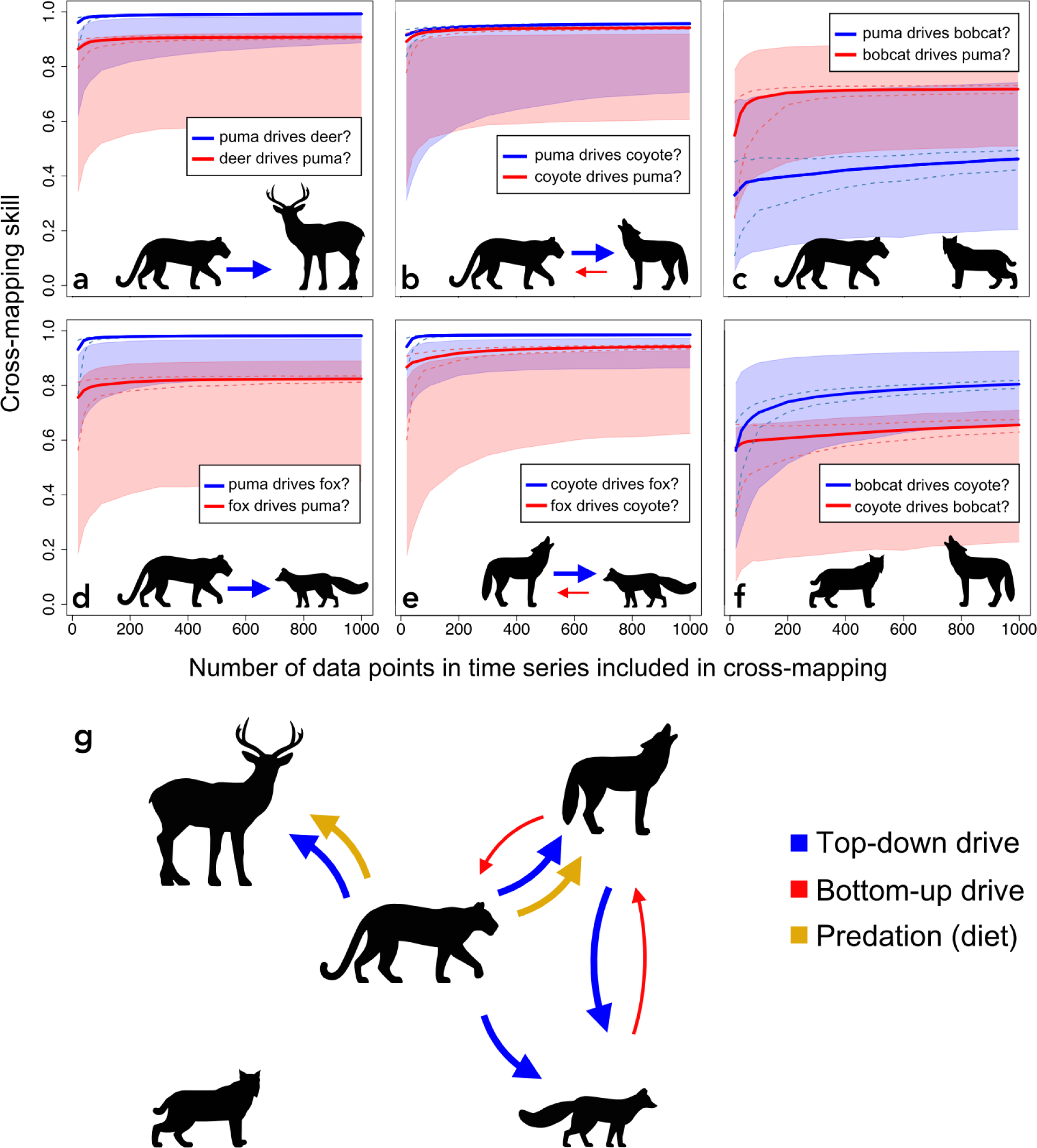
Inference of causal relationships between species. (a-f) CCM analyses of RAI from the 7-year dataset (see Figure S3 for the 5-year dataset). In general, top-down regulation (blue lines) has a higher cross-mapping skill than bottom-up regulation (red lines). Dashed lines represent the 2.5th and 97.5th quantiles of bootstrapped time series fragments. The number of data points refers to the length of the time series fragments used for cross-mapping. The cross-mapping skill is the Pearson’s correlation coefficient between observed and predicted values of the driver using the manifold constructed from the response variable. The shaded regions represent the 0th and 95th percentiles (95% upper one-sided bound) of the CCM null distributions (1000 runs of randomized time series). Arrows indicate the direction of causality based on significant CCM results (p < 0.05). Larger arrows indicate stronger drivers (higher cross-mapping skill). All cross-mappings showed significant convergence (Kendall’s test τ > 0 and p < 0.01). (g) Relationships between species based on CCM and diet analysis.

Mule deer DNA was present in all puma scat (Frequency of Occurrence; FOO=1), dominating the diet. Coyote was also found in puma’s diet (FOO=0.08). The diet of the coyote overlaps by 45% with the puma (See Table S3) and mostly consisted of rodents (Operational Taxonomic Unit; OTUs=4; FOO=0.09-0.64), deer (FOO=0.27), and lagomorphs (OTUs=2; FOO=0.09-0.36). Gray fox and bobcats mainly consumed small mammals (rodents and lagomorphs) and overlap substantially with the coyote (71 and 74% respectively), but not with the puma (14 and 19% respectively).

Finally, we found that multiple species changed their daily activity cycle after the increased abundance of puma (T3) compared to before (T1). Mule deer (Figure 3 and S7-8) became 42% less active at night, compensated by an increase in activity at dawn and before dusk. Coyotes became 27% more active during the day (Figure 3). Bobcat and puma activity remained largely unchanged and predominantly nocturnal (Figure S6 and S9).

**Figure 3.**
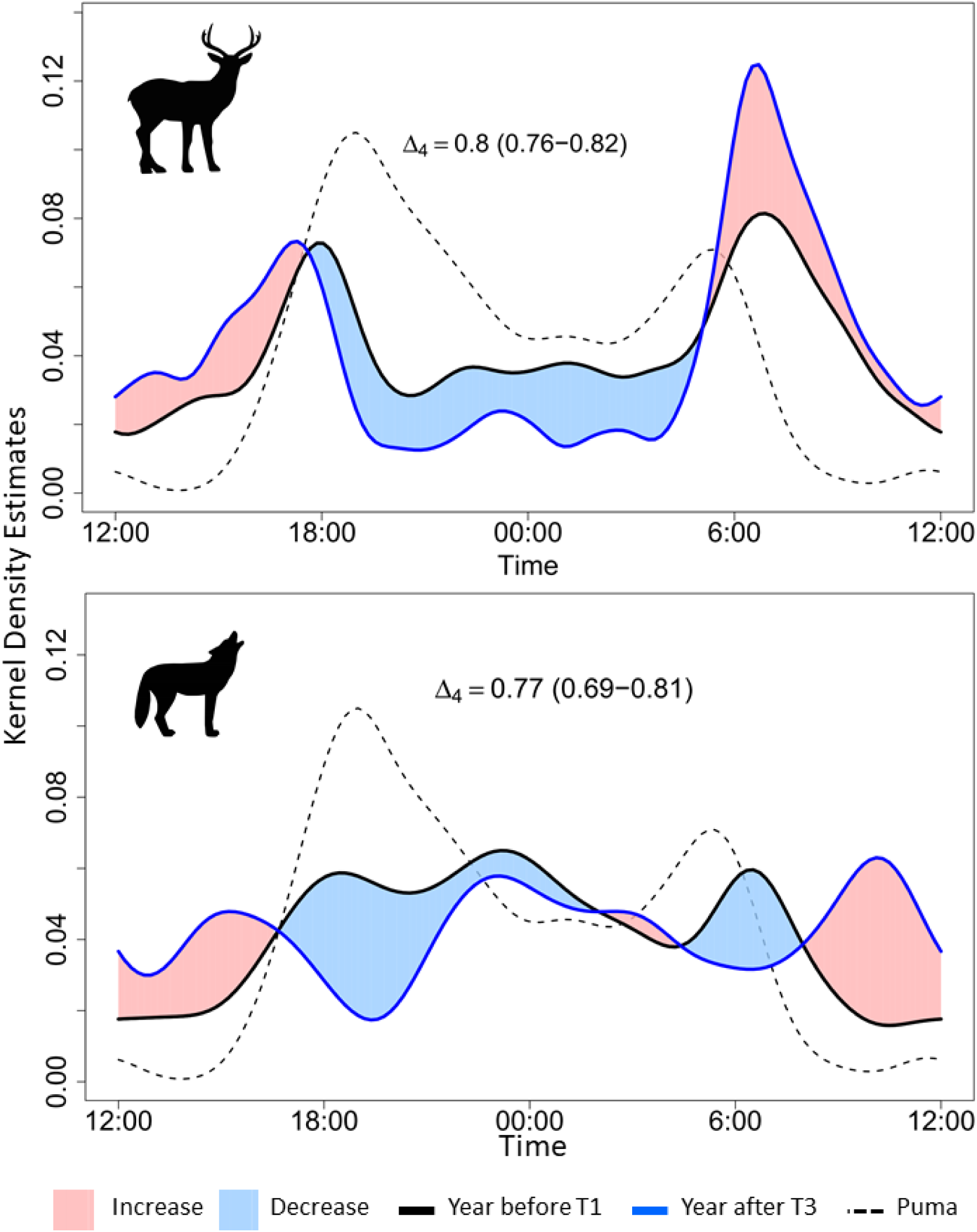
Kernel density estimates of daily activity patterns of mule deer (top) and coyote (bottom) between the year before T1 (return of resident pumas, black line) and the year after T3 (Coyote reach an equilibrium, blue line), from the 7-year dataset. Shaded areas correspond to increases or decreases in daily activity between these two years. Dashed line corresponds to the daily activity pattern of puma the year after T3. Overlap coefficient Δ_4_ between the two years varies between 0 (no overlap) and 1 (complete overlap). 95% confidence interval obtained with 1000 bootstraps is given in parentheses.

## DISCUSSION

Pumas natural return to JRBP in 2013 caused a trophic cascade that is strongly supported by multiple lines of evidence. First, time series of RAI and detection probability show a strong increase in pumas in just 18 months and an immediate, coincident decrease in mule deer and coyotes (59% and 86% decrease in RAI respectively in 33 months between T1 and T3). Coyote group sizes also declined, indicating they became more transient than resident^8^. By provoking the decline of coyotes, which were the prior dominant predator in the preserve, pumas have allowed smaller carnivores such as gray foxes to fill the canid niche^19–22^. Second, convergent cross-mapping validates that the pumas are exerting a top-down influence on this cascade. Third, fecal DNA analyses of pumas demonstrate that their primary prey is deer and occasionally coyotes. Fourth, dietary preference of deer, and even coyote, by pumas is corroborated by the ‘ecology of fear’^23^ we see in the divergent diurnal activity of deer and coyote following the rise of puma in our study: deer and coyote are less active during nightly periods of higher puma activity. The response of bobcats to pumas remains ambiguous as previously reported in the region^17,24^. However, unlike the other species, the bobcat results depend on which dataset we use. The RAI and CCM indicate a direct interaction between bobcats and pumas in the 5-years dataset, but not in the 7-year dataset, which we interpret as a difference in the particular placement of the newer cameras in areas preferred by bobcats. Importantly, bobcat diet almost exclusively contains small rodents and lagomorphs, suggesting no direct competition for food with the puma.

The trophic recovery may have other indirect effects on the ecosystem. Our results confirm that coyote infrequently consume large herbivores and favor smaller mammals^25^, while the puma diet is dominated by mule deer (as documented elsewhere in California^26^). This puma-mediated suppression of large herbivores may thus impact plant diversity and demography^27^. While we do not have data to support this effect here, the noticeable absence of browsing at sites where pumas are most frequently present could impact tree regeneration, which has been documented elsewhere to be induced by a landscape of fear^28^. Similarly, we noted the presence of seeds in all of the gray fox scat, which may play a large role in seed dispersal^29^, both in abundance and distribution as gray foxes become more common. Finally, by hunting mule deer, pumas generate an increasing number of carcasses, which are sources of food for carrion-dependent invertebrates^30^, smaller predators and scavenger birds such as turkey vultures^31^. Mule deer DNA found in the diet of all mesopredators could thus be explained by consumption of carcasses. Moreover, cameras deliberately set at deer carcasses observed this menagerie of scavengers, culminating with visits by turkey vultures, which have been increasing in the preserve (Figure S5).

The dense and permanent infrastructure of cameras traps presented here has allowed us to document a trophic cascade with an unprecedented level of detail. While this type of monitoring is not feasible everywhere, there are no technical barriers to its spread in suburban environments.

Most importantly, our study shows that small biological islands should not be abandoned in these highly fragmented landscapes dominated by humans. We show that trophic recovery in such landscapes is possible over a short period of time, provided the conditions favoring these large predators are met^11,17,32^. In this preserve, these conditions might include a limited public and vehicle access, low human density and being unused at night. In addition, the preserve is in close proximity to the Santa Cruz Mountains, which are largely protected from urbanization and have an abundance of pumas^15,17,31^. Finally, surrounding residential areas are of low density, typically unfenced, and replete with tree-lined drainages. These conditions seem to allow the puma to dominate the adaptable and synanthropic coyote, which has otherwise significantly expanded its distribution across North America as a result of the extirpation of larger predators^33,34^. As such, small suburban preserves are not only refuges for rare species^35^, they also support functional ecosystems where the top-down forcing of an apex predator can be realized. In the Anthropocene, these protected areas have a decisive role to play in stopping the erosion of biodiversity and, therefore, must be given immediate priority in conservation.

## METHODS

### Study area and camera traps

Jasper Ridge Biological Preserve (JRBP; Stanford, CA) covers a surface of 4.9 km^2^ in the vicinity of the Santa Cruz mountains. It is a partially fenced preserve, not accessible to the public and with limited usage of motor vehicles. Fourteen Buckeye Orion and 4 Buckeye X7D wireless camera traps were installed between 2009 and 2015 to serve multiple purposes and did not follow a probability-based sampling design. The initial setup served as a proof of concept for wireless system and was used to monitor for human trespassers as well as wildlife before pumas were first observed. The cameras were installed at strategic locations, usually trail intersections and sections of trail passing through geographic choke points to maximize detection of wildlife. Four of the cameras were placed specifically to serve as repeater cameras, which relayed the wireless signals around topographic obstacles. Fourteen of the cameras were equipped with solar panels while the remaining 4 were in locations with virtually no solar exposure and ran on batteries alone, which had to be changed every few months. The wireless cameras are managed via a computer-based software interface which allows the user to remotely configure and control the camera as well as view battery level and wireless signal quality. Camera status was monitored daily and battery health maximized by avoiding discharging the batteries below 50% whenever possible. As a result, all of the cameras have been continuously in operation with few and insignificant gaps in service. Automated scripting was used to copy the photos to a server located offsite for processing and backup. A custom web-based tagging interface was created and used by a group of volunteers to label species captured in each photo. The classified photos were then rechecked by at least one other person.

**Figure 3.**
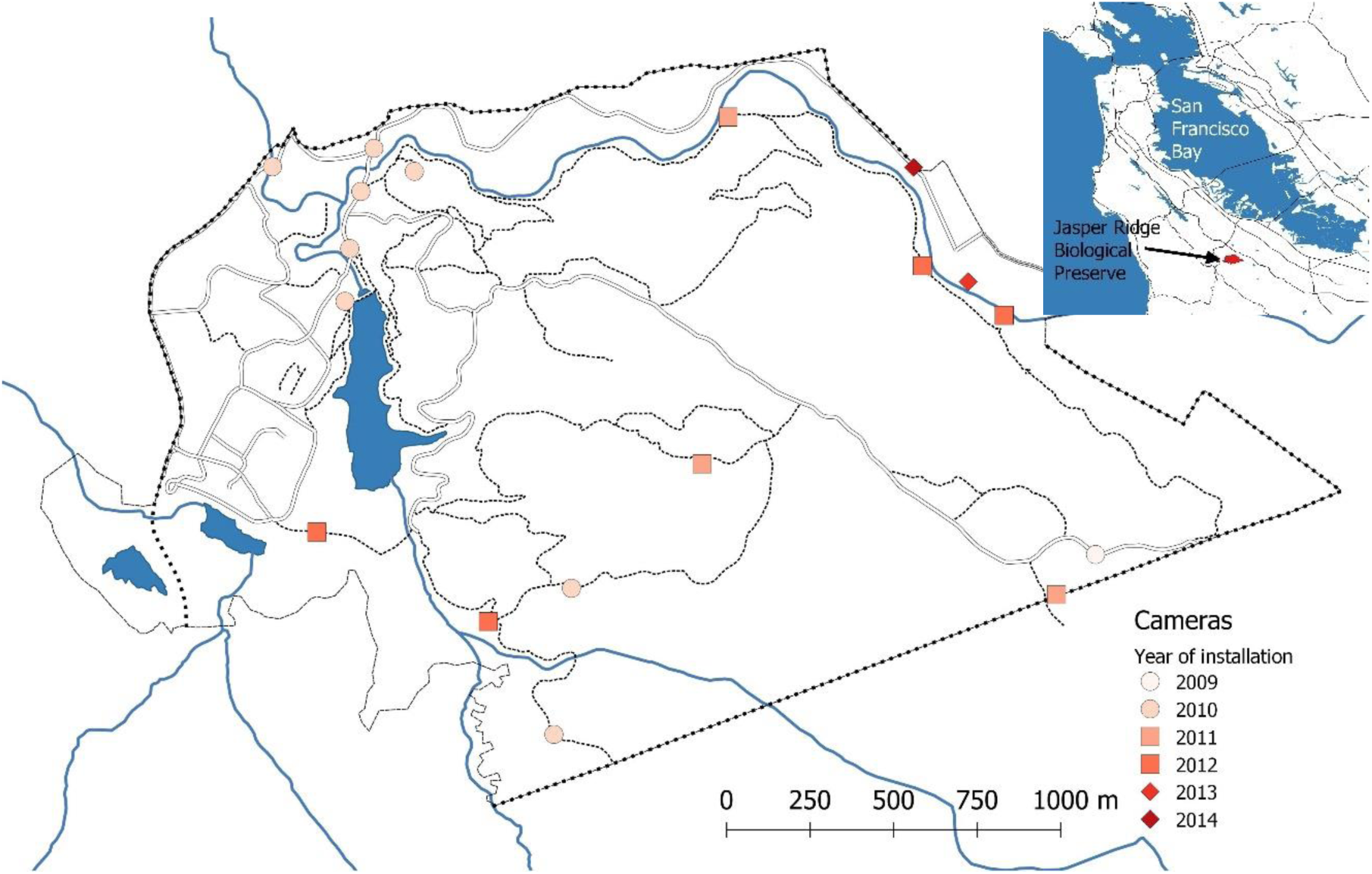
Map of Jasper Ridge Biological Preserve, Stanford, CA. Points corresponds to camera locations and are categorized by year of installation. Circular points correspond to the 7-year dataset while the combination of circular and squared points correspond to the 5-year dataset.

#### Filtering Camera-Trapping Data

Out of all species recorded, we focused on 11 of them and discarded flying birds, mammals with a mass smaller than 0.5 kg and species recorded less than 50 times over the whole survey. The species considered are: black-tailed jackrabbit (*Lepus californicus*), bobcat (*Lynx rufus*), brush rabbit (*Sylvilagus bachmani*), coyote (*Canis latrans*), gray fox (*Urocyon cin ereoargenteus*), mule deer (*Odocoileus hemionus*), puma (*Puma concolor*), raccoon (*Procyon lotor*), striped skunk (*Mephitis mephitis*), Virginia opossum (*Didelphis virginiana*) and wild turkey (*Meleagris gallopavo*).

Photographic events of the same species were considered independent if they occurred more than 30 min after the previous photo of that species at the same camera^17^. Pictures containing multiple individuals were also treated as one event. Because camera traps were installed progressively over time, we decided to define 2 datasets for our analysis. The first contains the records of 9 cameras from October 26, 2010 until June 30, 2017, referred to as the 7-year dataset. The second contains the records of 16 cameras from October 2, 2012 until June 30, 2017, referred to as the 5-year dataset.

#### Species Relative Abundance and Detectability

We calculated the relative abundance index (RAI) as the number of events per camera trap per year. RAI was thus calculated for each day with a 1 year moving window for both the 7-year and 5-year datasets. In addition, we calculated the detection probability using the R package unmarked ^36^. Detection/No-detection per camera were calculated in periods of 1 week for both datasets. Detection probability was then calculated for periods of 1 year (52 weeks) with a step of one week.

### Empirical Dynamic Modeling

We used the RAI time series to perform empirical dynamic modeling (EDM); an approach that detects putative causal relationships in nonlinear systems^18^. First, we standardized the time series to zero mean and unit variance for unbiased comparability between species abundance. Next, we used an EDM method called convergent cross-mapping (CCM)^18^ with simplex projection^37^ to infer causal relationships between species. CCM cannot, however, distinguish between the different types of relationships, *e.g*., competition *versus* predator-prey dynamics, or how abundance is mediated (*e.g*., through birth-death processes, change in diel activity, or migration). However, we also conducted diet and behavioral analyses to address this gap.

The CCM method uses time series of different variables (*e.g*., different species RAI) to reconstruct dynamics of a system by constructing a state-space manifold. Here, the manifold represents the different states of the ecosystem of the species included in this study. This method uses the property of *Takens’ Theorem*^38^, which states that a manifold, *M*, representing a system can also be reconstructed using just one of the variables (*e.g*., puma RAI) lagged against itself (*e.g*., *X*(*t*), *X*(*t*–τ), *X*(*t*–2τ) for variable *X* and time lag *τ*). This creates a univariate shadow manifold *M*_*X*_ that preserves the properties of the original manifold *M*. CCM detects causal relationships between variables *X* and *Y* by comparing local points *x*(*t*) and *y*(*t*) on shadow manifolds *M*_*X*_ and *M*_*Y*_, respectively ^18^. For example, if puma is driving deer RAI, either through predation or by changing deer behavior, then information about puma RAI will be embedded in the dynamics of deer RAI, such that the shadow manifold *M*_*deer*_ can reconstruct past values of puma RAI.

The first step of EDM was to construct a univariate shadow manifold from each individual time series. The optimal number of lagged times series plus the original time series—i.e., the embedding dimension *E* used to construct the manifold—was obtained by performing a nearest-neighbor prediction method called simplex projection^37^. The *E* that generated the highest prediction skill ρ (Pearson’s correlation coefficient between observed and predicted values using simplex projection), was chosen for the reconstruction of shadow manifolds to be compared (cross-mapped) when performing CCM. The cross-mapping between the dynamics of a putative driver (*e.g*., puma abundance) and the dynamics of a putative response variable (*e.g*., deer abundance) is, again, performed using simplex projection. If there is a causal signal in the data, then the longer the time series, the denser the shadow manifold becomes, and the shorter the distance between nearest-neighbors becomes, leading to higher prediction skill. This phenomenon, called *convergence*, is an essential criterion for CCM to detect causal relationships.

We used a null model to assess the significance of the CCM results for causality between a pair of variables (*i.e*., a pair of species RAI). For the null model we created several randomized surrogate time series of the putative driver variable for the cross-mapping, losing any signal of causality if present in the original time series. To account for spurious predictability just based on neighboring time-dependence (*i.e*., serial correlation) we used a strict null model that conserved any autocorrelation in each surrogate time series by Fourier-transforming the time series and only randomizing the phases before re-transforming the time series back to its original form. This is known as the Ebisuzaki method^39^. We obtained a null distribution with 95% confidence intervals from 1000 surrogates with randomized Fourier phases, and compared its CCM model performance with the true time series using a right-tailed z-test to obtain the p-value^40^. The variance (95% confidence intervals) of the CCM performance with the true time series was obtained from 1000 bootstraps of different time series lengths from randomized time series locations. In addition, to test the significance of convergence, we used the Kendall’s test^41^, which tests whether the cross-mapping skill ρ is significantly higher when using the whole time series compared to just one time point (if the statistic τ > 0 and p < 0.01 then convergence is significant). All EDM analyses were performed using the rEDM package in R^42^.

### Fecal Sample Collection & Preservation

Fecal samples were collected from October 2017 – April 2018 covering 32 paths (trails 17 km; roads 7 km) within JRBP. In total, over 175 km of trails were traversed over 23 collection days. Whole scat samples were collected in sterile bags and using gloves to avoid contamination. All samples were stored at −20C until DNA extraction. Over the wet and dry season, 157 predator scat samples were collected (puma=15, coyote=11, bobcat=49, grey fox=82).

#### DNA extraction, amplification and sequencing

Scat samples were thawed, homogenized and processed (∼0.2 mg) utilizing Zymo Quick-DNA Fecal/Soil Miniprep Kit^43^. Samples were processed in small batches (∼ 14) with an extraction blank to monitor for potential cross-contamination in the laboratory. The eluted DNA was quantified using Nanodrop 2000 (Thermo Fisher Scientific Inc).

Metabarcoding primers for the 12S mtDNA were selected that amplify DNA from a wide range of mammal species that are well represented in public databases. The MiMammal-U primers were used to amplify mammals specifically and modified with the Illumina adaptor preceding the target primers and separated by 6-N spacers^44^. The PCR comprised 20 µl: 10 µl of GoTaq® Colorless Master Mix (400μM dATP, 400μM dGTP, 400μM dCTP, 400μM dTTP and 3mM MgCl2), 1 μL of each primer (5mM), 4 μl of DNA template and 4 μl of H_2_0. Cycling conditions used initial denaturing at 95°C for 10 min, followed by 35 cycles of denaturing at 95°C for 30 s, annealing at 60°C for 30 s and extension at 72°C for 10 s. After visualization on a gel, PCR amplicons were purified using the QIAquick PCR Purification Kit (Qiagen).

For indexing, appropriate Illumina barcodes were ligated to each sample. The index PCR was performed as 20 µl reaction: 10µl of Amplitaq Gold reactions (with 2.5 mM MgCl2,200 lM each dNTP, 0.1 mg/mL BSA, 4% DMSO) 1.2 µl (of each primer), 1.6 μl of DNA amplicons and 6µl of H_2_0. Cycling conditions used initial denaturing at 95°C for 10 min, followed by 15 cycles of denaturing at 95°C for 30 s, annealing at 50°C for 30 s and extension at 72°C for 10 s.

The indexed second PCR products (n=118) were quantified and assessed for quality control and quantifying amplicon DNA yields using the Fragment Analyzer, normalized to equimolar concentrations and pooled together before purification using QIAquick PCR Purification Kit (Qiagen).

Sequencing was performed on a Miseq platform using the Reagent Kit Nano v3 for 2 × 300 bp PE (Illumina, San Diego, CA, USA) and run at Stanford University PAN Facility. A 30% PhiX DNA spike-in control was added to improve the data quality of low diversity samples such as single PCR amplicons used in this study. Samples were pooled with other projects on 2 Miseq runs, generating a total of 25,370,906 reads, or on average 256,271 reads per samples.

##### DNA metabarcode demultiplexing, quality control, and species identification

We used the software packages Obitools^45^ and R (R Core Development Team 2013) for demultiplexing and quality control. Each sequence was assigned to its sample of origin based on exact matches to both multiplex identifier (MID) tags. Sequences were paired with Obitools *illuminapairedend* and aligned sequences with a score <40 were discarded. Forward and reverse adapters were then removed in Cutadapt^46^. After assignment of sequences to their corresponding samples, we used *obiuniq* to dereplicate reads into unique sequences, eliminated potential PCR and sequencing errors with *obiclean*, and kept only sequences occurring at least 10 times.

Taxonomic assignment of sequences was done against a custom reference database. First, we downloaded all standard mammal, human, mouse and vertebrate sequences from embl (http://ftp.ebi.ac.uk/pub/databases/embl/release/std/) and converted recovered file to ecoPCR format. EcoPCR was then used to simulate an in-silico PCR, using the Mimammal-U primers and maximum 3 mismatches. *Ecotag* was then used to identify dietary sequence, while inspecting and revising taxonomic assignments to ensure validity. Sequences with poor matches to reference database (<95%) were removed. After quality control, our final data consisted of 99 samples for the diet analysis (puma=13, coyote=11, bobcat=30, grey fox=45).

Diet composition was quantified using Sequence Occurrence (*i.e*., presence/absence) which when averaged across all samples yields relative frequency of occurrence (FOO) and the mean sequence Relative Read Abundance (RRA) range defined as the proportion of unique Illumina sequence reads in a sample divided by the final (*i.e*., after quality control & removal of host species reads) number of sequence reads in that sample^43^.

We used Pianka’s adaptation of the niche overlap (*Ojk*) metric to determine diet overlap among all pairs of target carnivores^47^:

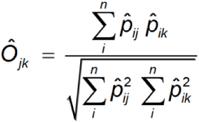

Whereas *pij* is the proportion of prey species *i* in carnivore species *j diet, pik* is the proportion prey species *i* in carnivore species *k diet, n* = Total number of available prey species and *Ojk* = 0 represents no overlap, whereas a value of *Ojk* = 1 represents complete overlap.

### Daily Activity Cycle

We looked for changes in daily activity cycle over time by measuring the overlap of activity between species (predator-prey). Because daily activity cycle of mammals depends largely on daylight rather than time of day, we considered the seasonal patterns in sunlight. To do so, we standardized all recording times in a standard day where sunrise, solar noon, sunset and solar midnight are set as 6am, 12pm, 6pm and 12am respectively. We obtained time of sun cycle for Stanford, CA, from the Astronomical Applications Department of the U.S. Naval Observatory (http://aa.usno.navy.mil/data/docs/RS_OneYear.php). We then rescaled the times of pictures recording into the standardized day depending on the day of observation. Our standardized daily activity cycle is thus representative of the control of daylight on animals’ activity. Next, we used the R package *overlap*^48^ to plot patterns of daily activity cycle. The overlap varies between 0 (no overlap of time of activity) and 1 (complete overlap of time of activity). Confidence intervals of 95% for the overlap were estimated with 1000 bootstraps. First, we looked for changes in daily activity cycle for the same species between the year before the cascade (T1) and the year after (T3). Second, we looked for changes of daily activity cycle within a species over time. For each species, we thus used the first 12 months to define a reference year and then compared it with each successive year, with a step of 1 month. Finally, we compared the daily activity cycle of predators and their prey. In some cases, there were not enough observations per species per year to produce an accurate representation of their daily activity cycle and we decided not to include them in the results. In addition, we considered that a minimum of 50 observations were necessary to use Δ_4_, and resorted to Δ_1_ otherwise^49^.

## ACKNOWLEDGMENTS

We would like to thank all the Jasper Ridge Biological Preserve docent volunteers who helped to identify the animals on pictures. We would also like to thank Alina Isakova for her help during the diet analysis, Fred Boyer for helping us troubleshooting Obitools, and Ethan Deyle for helpful guidance on empirical dynamic modeling. We also thank Rodolfo Dirzo for initiating the camera-trapping project at Jasper Ridge Biological Preserve.

K.L. was funded by an Early Postdoc Mobility grant from the Swiss National Science Foundation (P2ELP3_175075) and JRBP Kennedy Fund. N.N. was funded by The Bing Fellowship in Honor of Paul Ehrlich. J.M. was funded by the Philippe Cohen Graduate Research Fellowship. The Jasper Ridge wireless camera traps and supporting infrastructure were paid for by National Science Foundation (grant number 0934210).

## Contributions

K.L. performed the camera trap analyses, the bioinformatic part of the DNA metabarcoding, and wrote the initial draft. J.M. performed the fecal diet analysis. T.H. maintained the camera traps and curated the database. N.N. performed the empirical dynamic modeling and created the artwork in Figure 2. E.A.H. designed and supervised the project. All authors contributed to the writing of the paper.

