## Supplementary tables and figures for "Return of an apex predator to a suburban preserve triggers a rapid trophic cascade"

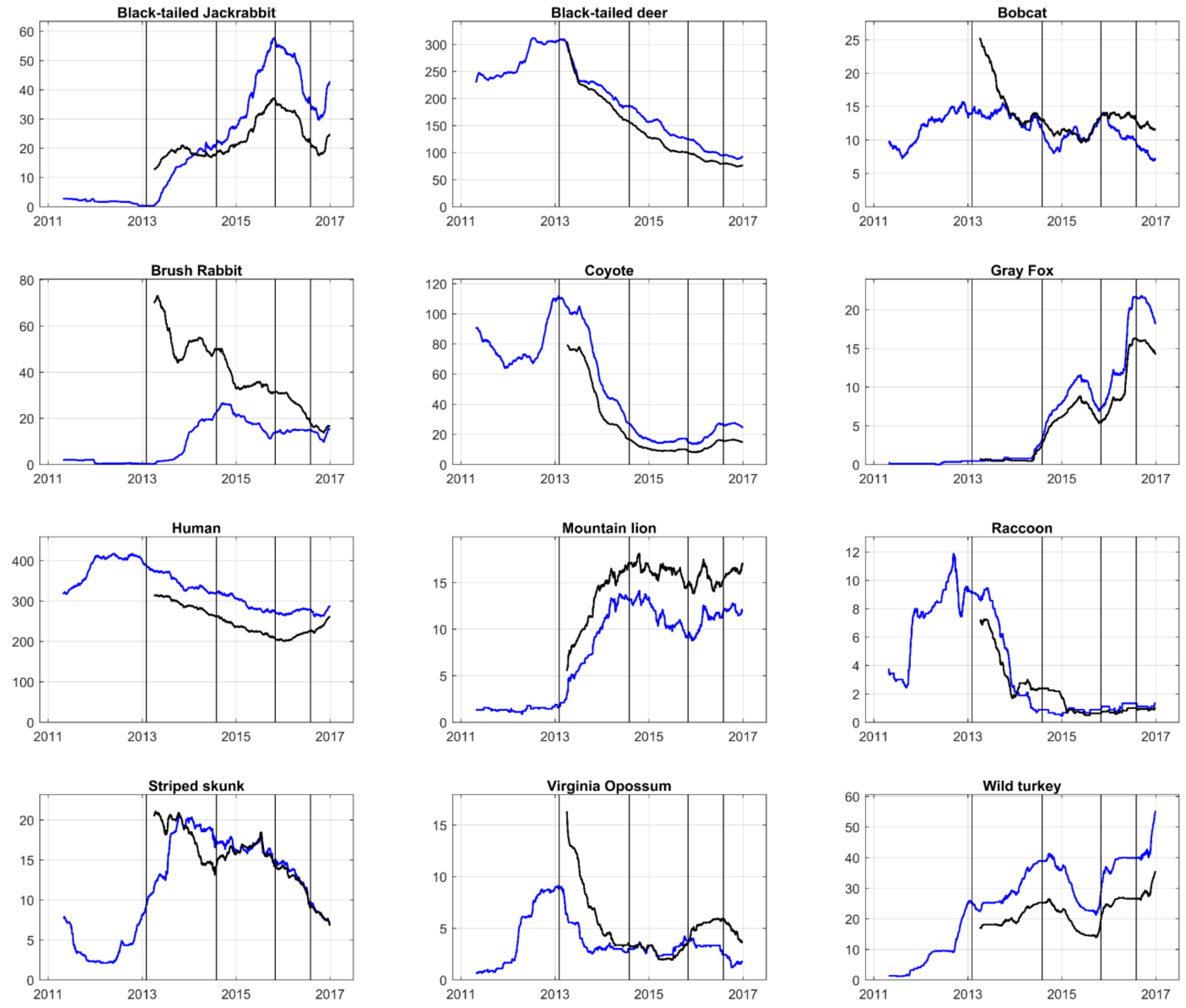


Figure S1. Relative Abundance Index (RAI) of the 12 species included in the 7-year (blue line) and 5-year (black line) datasets using 365 days moving window (step = 1 day) and divided by the number of cameras. Dates of time points: T1: 01-02-2013; T2: 01-08-2014; T3: 01-11-2015; T4: 01-08-2016.

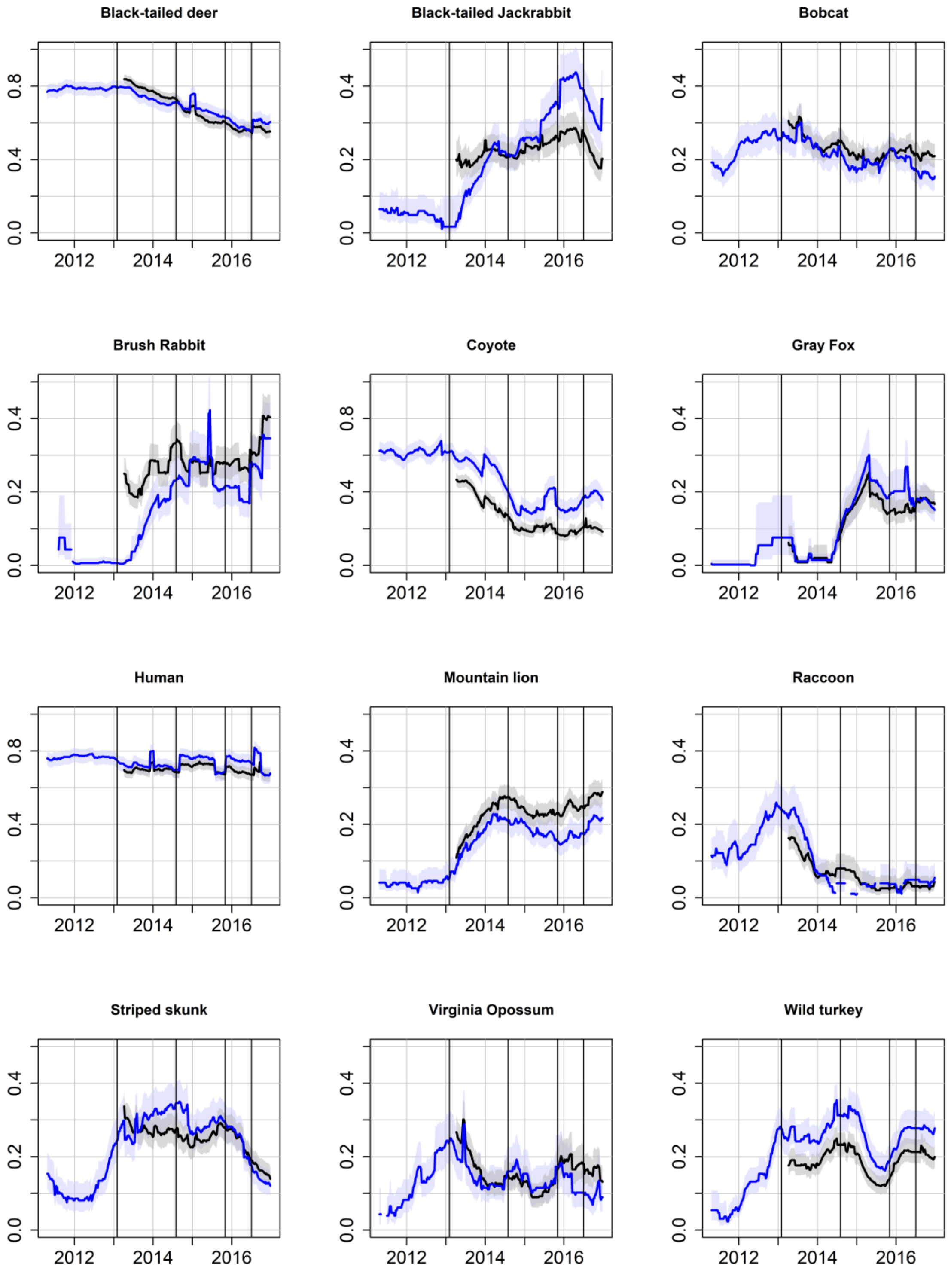


Figure S2. Detection probability of the 12 species included in the 7-year (blue line) and 5-year (black line) datasets using 52 collections of 1 week. Dates of time points: T1: 01-02-2013; T2: 01-08-2014; T3: 01-11-2015; T4: 01-08-2016.

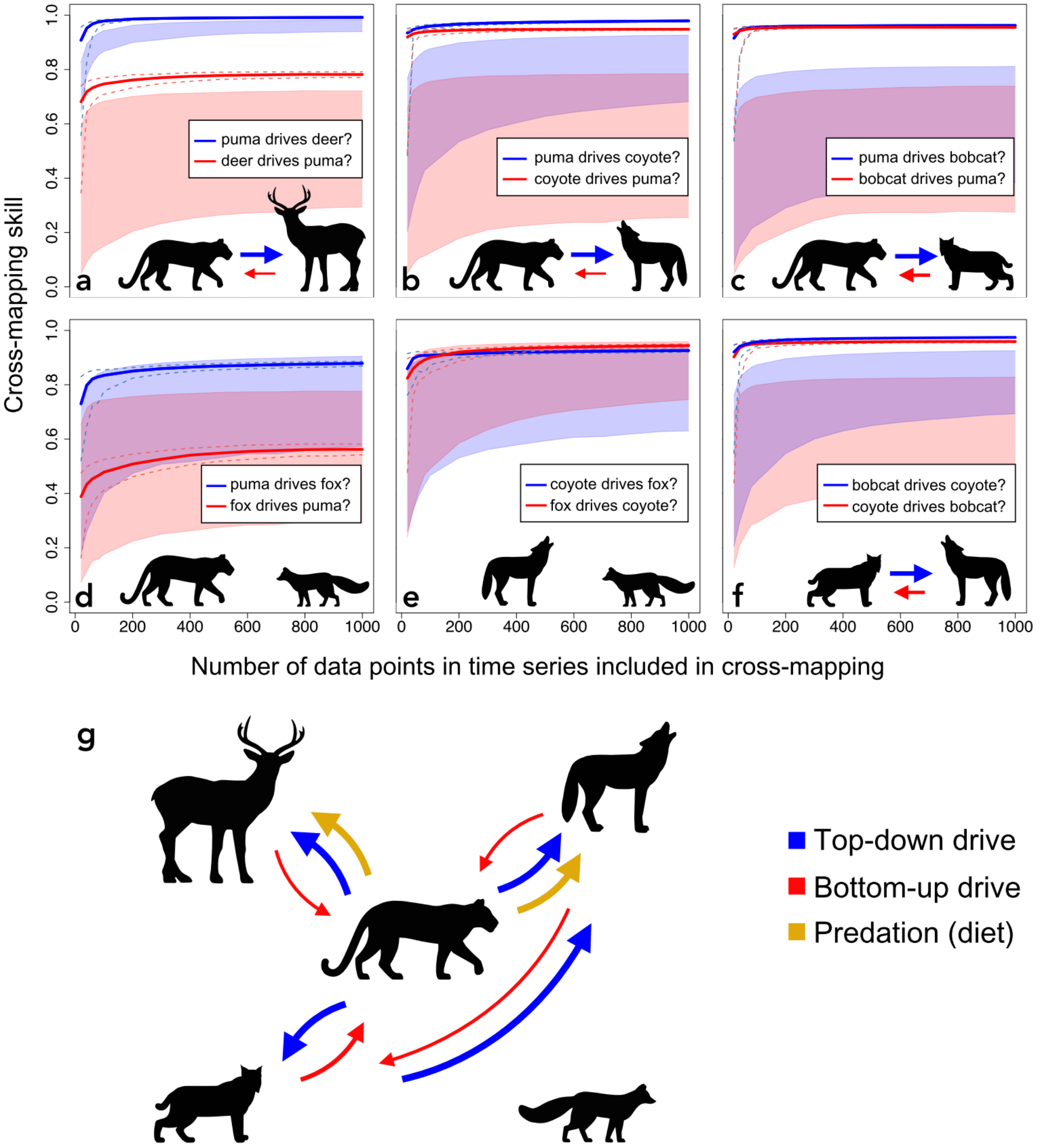


Figure S3. Inference of causal relationships between species. (a-f) CCM analyses of RAI from the 5-year dataset. In general, top-down regulation (blue lines) has a higher cross-mapping skill than bottom-up regulation (red lines). Dashed lines represent the 2.5th and 97.5th quantiles of bootstrapped time series fragments. The number of data points refers to the length of the time series fragments used for cross-mapping. The cross-mapping skill is the Pearson's correlation coefficient between observed and predicted values of the driver using the manifold constructed from the response variable. The shaded regions represent the 0th and 95th percentiles (95% one-sided upper bound) of the CCM null distributions (1000 runs of randomized time series). Arrows indicate the direction of causality based on significant CCM results ($p < 0.05$). Larger arrows indicate stronger drivers (higher cross-mapping skill). All cross-mappings showed significant convergence (Kendall's test $\tau > 0$ and $p < 0.01$, except subfigure f had $p = 0.02$). (g) Relationships between species based on CCM and diet analysis.

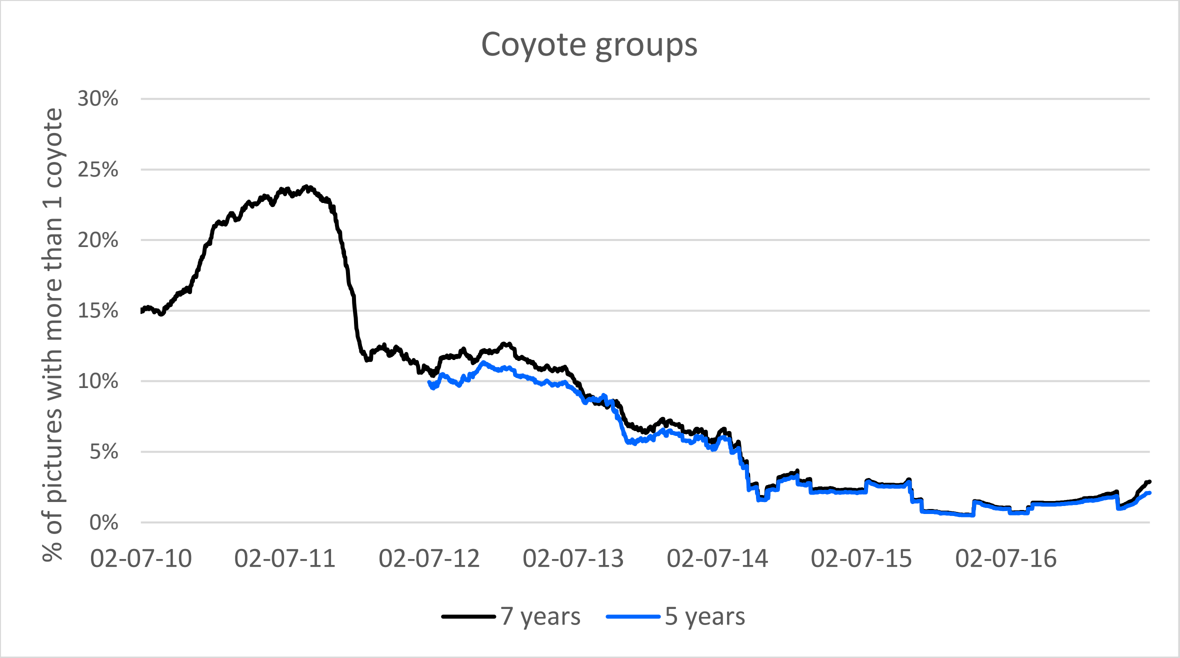


Figure S4. Proportion of coyote events involving more than 1 individual. Based on events from the 7 years (black) and 5 years (blue) datasets. Proportion is calculated per year (365 days moving window) with a step of 1 day.

Table S1. Human use per camera. Average density of humans per day at each camera using the 5y dataset.

| Camera | Average Density (People/Day) |
|---|---|
| F1 | 1.811 |
| B12 | 1.997 |
| B6 | 2.52 |
| D1 | 0.371 |
| B3 | 0.337 |
| B13 | 0.698 |
| B8 | 0.914 |
| B11 | 0.378 |
| B5 | 0.284 |
| C2 | 0.329 |
| A2 | 1.082 |
| C3 | 0.021 |
| B2 | 0.008 |
| E1 | 0.113 |
| E3 | 0.068 |
| B4 | 0.006 |

Table S2. Diet overlap between predators, as calculated with Pianka's niche overlap index.

| Niche overlap - Pianka's index | | | | | |
|---|---|---|---|---|---|
| **Puma - Coyote** | **Puma - Bobcat** | **Puma - Fox** | **Coyote - Bobcat** | **Coyote - Fox** | **Bobcat - Fox** |
| **0.45** | **0.19** | **0.14** | **0.74** | **0.71** | **0.90** |

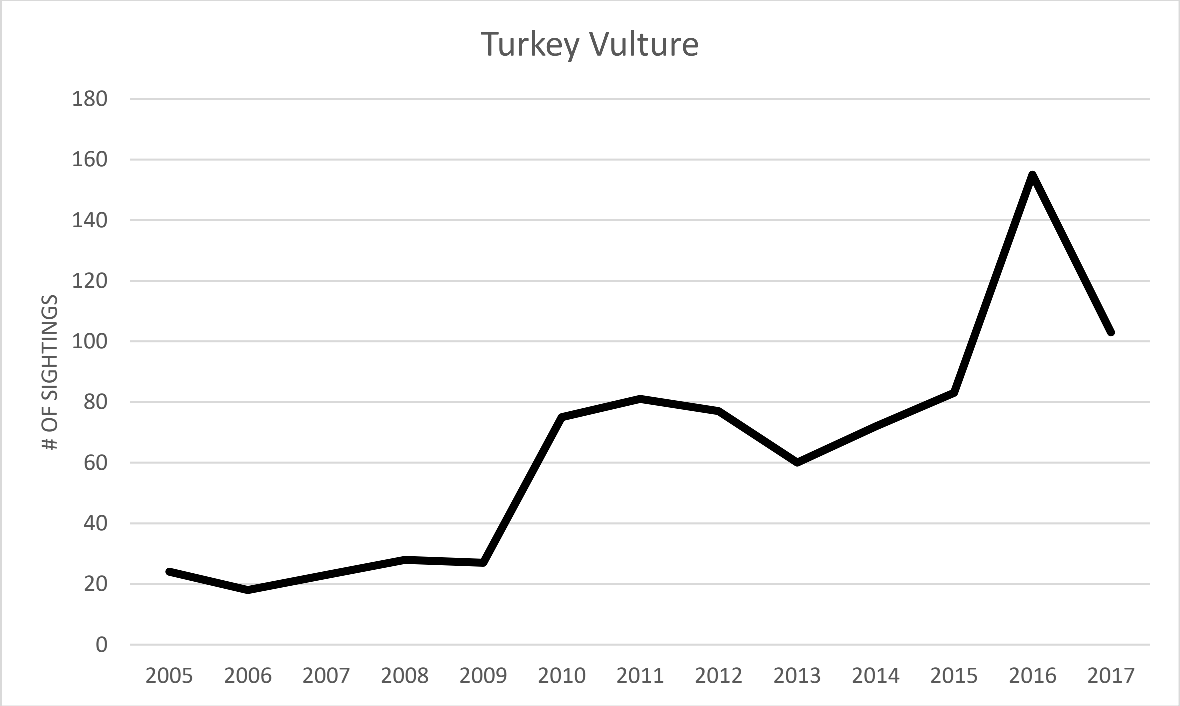


Figure S5. Transect survey data for Turkey vultures. Collected monthly by teams of expert birders along 6 fixed trail routes at Jasper Ridge. http://jrbp.stanford.edu/research/projects/jasper-ridge-bird-monitoring-program.

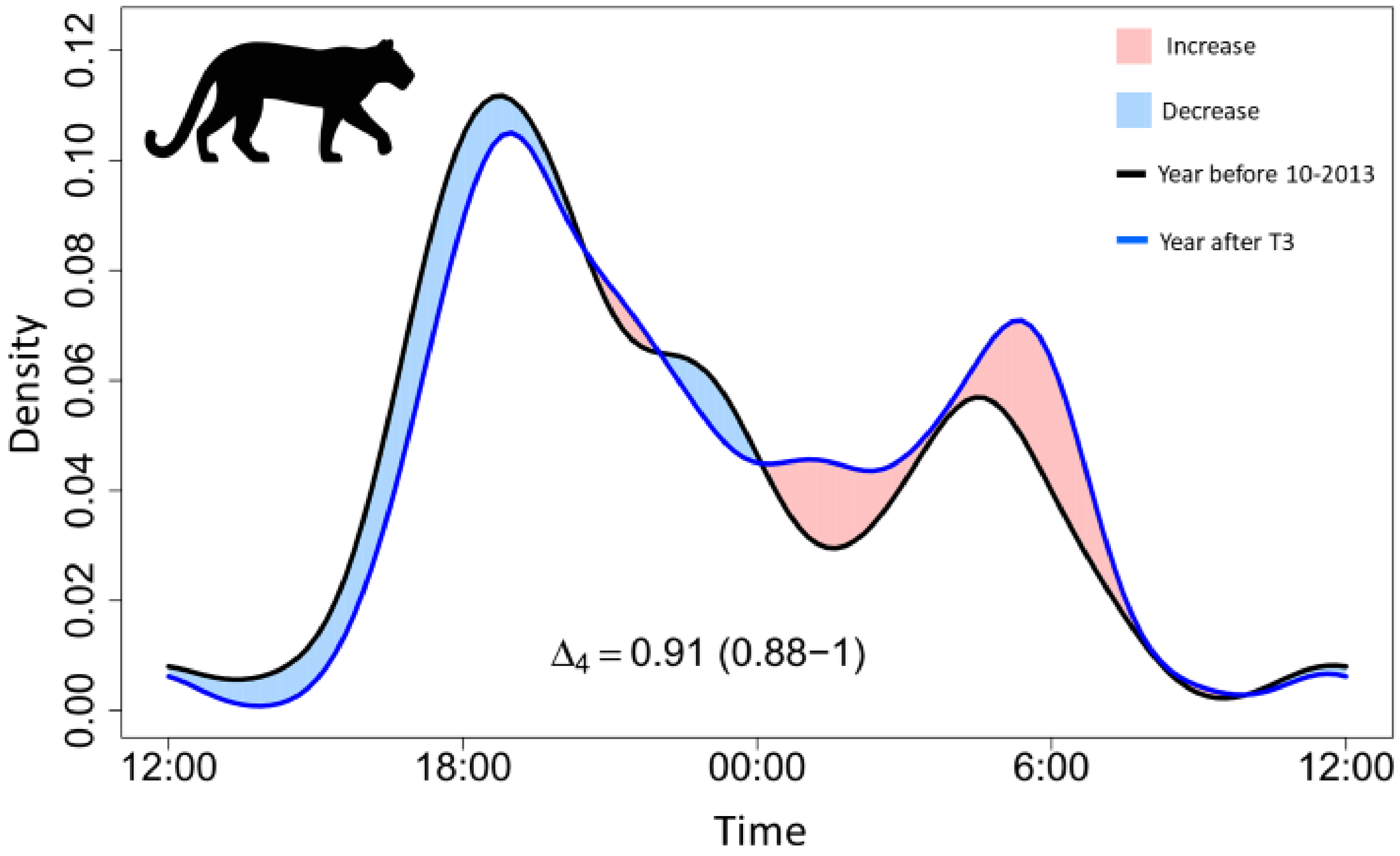


Figure S6. Overlap of daily activity cycle for pumas between the first year of the 5-year dataset (black line) and the year after T3 (blue line). Overlap coefficient $\Delta_4$ varies between 0 (no overlap) and 1 (complete overlap). 95% confidence interval obtained with 1000 bootstraps is given in brackets.

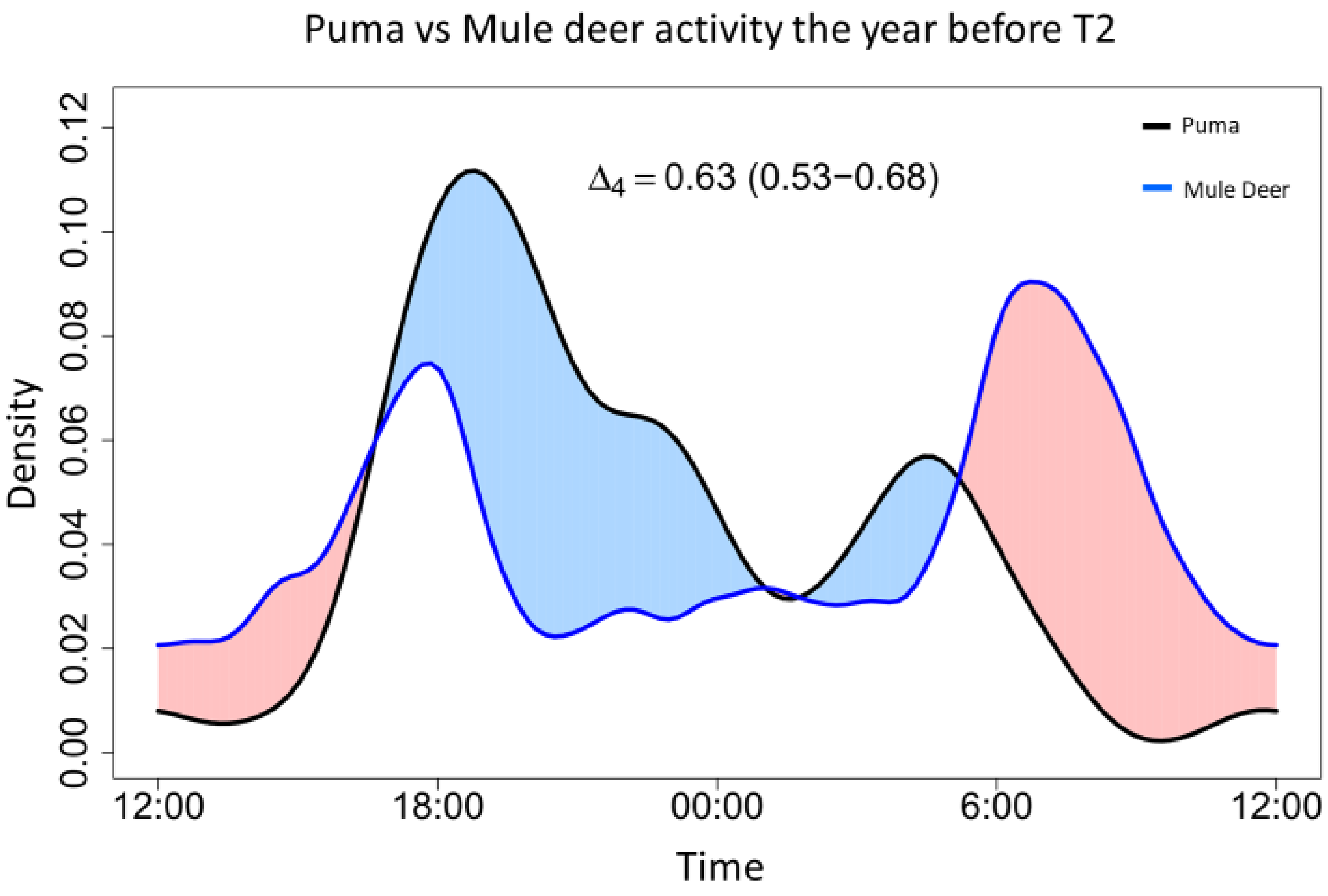


Figure S7. Overlap of daily activity cycle between pumas (black line) and deer (blue line) the year before T2 using the 5-year dataset. Overlap coefficient $\Delta_4$ varies between 0 (no overlap) and 1 (complete overlap). 95% confidence interval obtained with 1000 bootstraps is given in brackets.

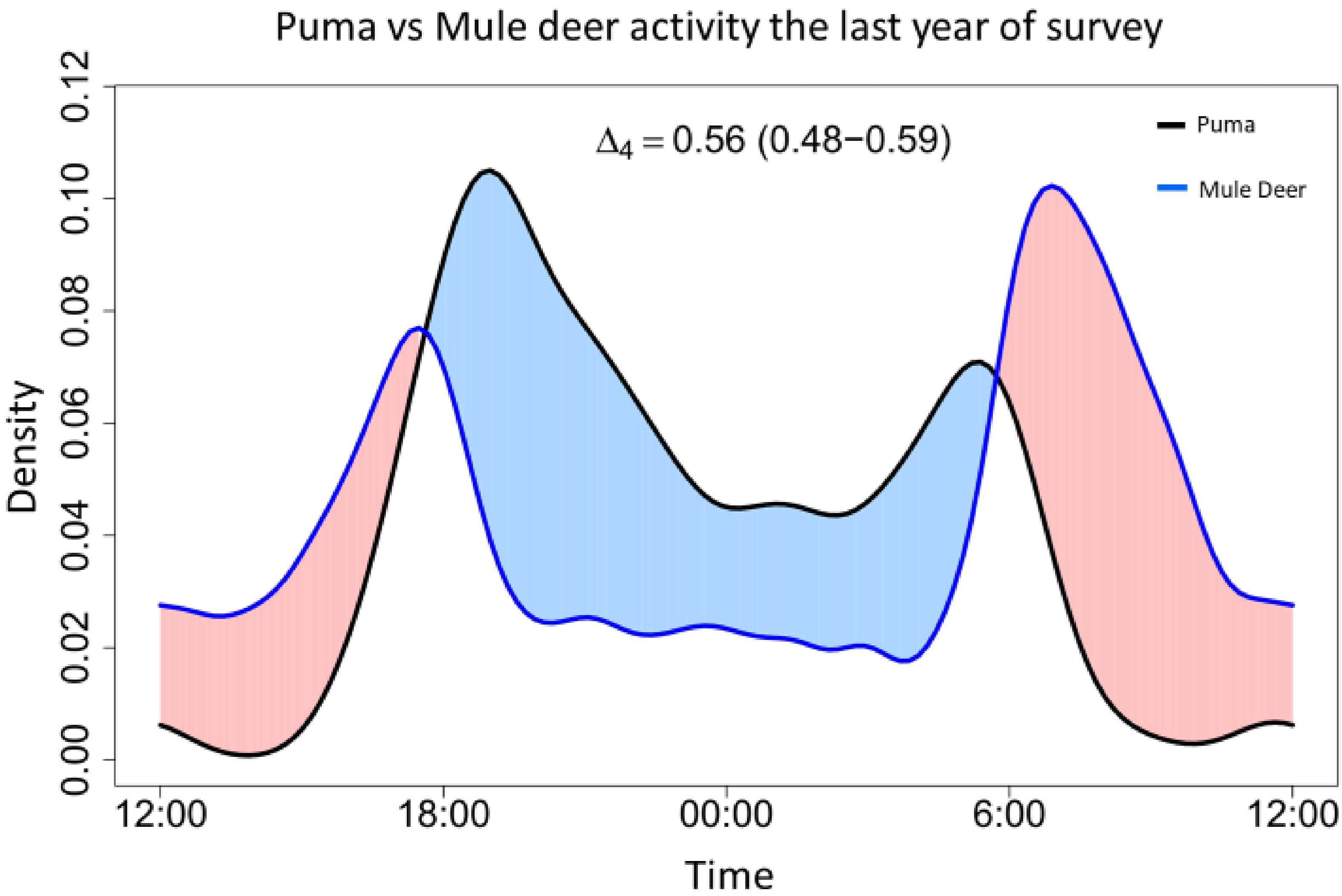


Figure S8. Overlap of daily activity cycle between pumas (black line) and deer (blue line) the last year of survey using the 5-year dataset. Overlap coefficient $\Delta_4$ varies between 0 (no overlap) and 1 (complete overlap). 95% confidence interval obtained with 1000 bootstraps is given in brackets.

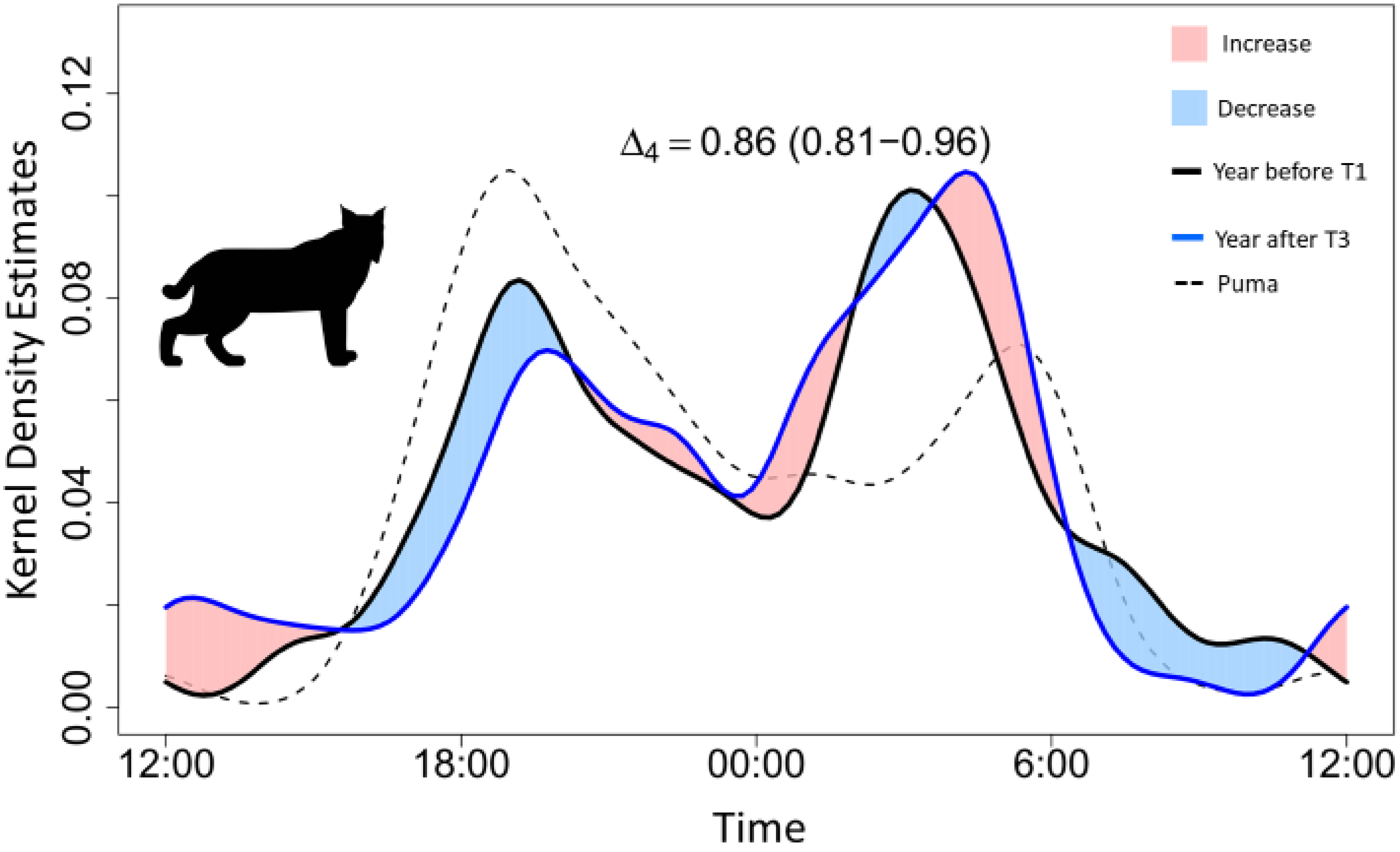


Figure S9. Overlap of daily activity cycle for bobcats between the year before T1 (black line) and the year after T3 (blue line), using the 7-year dataset. Dashed line corresponds to the daily activity pattern of puma the year after T3. Overlap coefficient $\Delta_4$ varies between 0 (no overlap) and 1 (complete overlap). 95% confidence interval obtained with 1000 bootstraps is given in brackets.
